# But, what are the cells doing? Image Analysis pipeline to follow single cells in the zebrafish embryo

**DOI:** 10.1101/2023.06.01.543221

**Authors:** Arianne Bercowsky-Rama, Olivier F. Venzin, Laurel A. Rohde, Nicolas Chiaruttini, Andrew C. Oates

**Affiliations:** Institute of Bioengineering, École Polytechnique Fédérale de Lausanne; Lausanne, CH; BioImaging and Optics Core Facility, École Polytechnique Fédérale de Lausanne; Lausanne, CH

## Abstract

Microscopy has rapidly evolved at pace with live markers, enabling ever higher spatiotemporal resolution of multicellular dynamics within larger fields of view. Consequently, we are now in the era of widespread production of terabyte (TB)-sized time-lapse movies of experimental model systems, including developing embryos and organoids. Working with these large datasets has presented a new set of hurdles, particularly due to the lack of standardized open-source pipelines for acquiring, handling and analyzing the data. Moreover, although long-term tracking of a cell throughout an entire process, for example vertebrate organogenesis, is key to revealing the underlying cellular dynamics, this has proven largely elusive. To specifically address the question “But, what are the cells doing?”, we created an image analysis pipeline optimized to track single cells in light-sheet acquired datasets (1 TB sized time-lapse, 8h of imaging, 30 min gene expression cycle, cell movement speed (1µm /1 minute), 200-400 µm tissue depth). Our modular pipeline optimizes and connects the following: image acquisition parameters to improve tracking feasibility; hardware specifications; data handling and compression tools; pre-processing steps; state-of-the-art cell tracking tools (Mastodon, MaMuT) and a novel open-source/ python-based tool (Paleontologist) to analyze and visualize spatiotemporal dynamics of the tracked cells. Importantly, our pipeline is adaptable to a variety of experimental systems and accessible to researchers regardless of expertise in coding and image analysis.

**One-sentence Summary:** User-friendly cell-tracking pipeline that connects image acquisition in multicellular systems through to data analysis of cellular dynamics.

## Introduction

Live imaging of multicellular systems to uncover tissue and cellular spatiotemporal dynamics has become an important line of research in many labs (Attardi et al., 2018; McDole et al., 2018; Shah et al., 2019). We have previously applied this approach to describe the cellular-level dynamics underlying the segmentation clock wave pattern in the developing zebrafish embryo (Rohde et al., 2021; Soroldoni et al., 2014). Here we detail the pipeline we created to facilitate imaging, cell-tracking, and data analysis of rapid gene expression oscillatory dynamics (30-min cycles) and cell movements of individual cells throughout the hours-long timeframe of vertebrate body segmentation in the zebrafish embryo. This pipeline is modular and adaptable to other systems, including organoids, in which researchers wish to track spot-like structures.

When imaging a tissue at cellular resolution, the ultimate goal is often to quantify the spatiotemporal dynamics of individual cells. This necessitates tracking the cells over time. There are two main approaches to cell tracking, the first of which is *in toto* cell tracking, such as performed by McDole et al., 2018, and Shah et al., 2019, in mouse and zebrafish embryos, respectively. These *in toto* approaches relied on automatic algorithms, including TGMM (Tracking with gaussian mixture model, Amat et al., 2014), to generate the cell tracks in the order of many thousands, a scale that renders manual curation unrealistic. Automated tracking accuracy decays exponentially over trajectory length, thus limiting analysis to short tracks as in Shah et al., 2019 (cell tracks of 10 frames, 20 minutes), or requiring custom statistical analysis to infer the dynamics as in McDole et al. 2018 (less than 30 time points over a 2-hour period). The second cell-tracking approach relies on manual or semi-automatic cell tracking (Delaune et al., 2012; Shih et al., 2015; Rohde et al., 2021), in which the user selects cells within a region of interest then manually curates the tracks. Although the number of tracks obtained is lower, in the order of many hundreds, this approach produces reliable trajectories that run considerably longer (300 frames, 450 min, Rohde et al., 2021 using Mastodon). Selection of one of these two approaches will depend on the question being asked and the accuracy required to answer it. Here, our pipeline takes a semi-automatic tracking approach, but includes optimized parameters for both imaging and processing steps to reduce the burden of manual curation.

Despite examples of successful cell tracking and analysis at various scales and timeframes, it remains out-of-reach for many labs due to lack of expertise. A diverse set of skills is required across the many steps of the process, including the following: 1) preparing and mounting live samples (Kleinhans and Lecaudey, 2019; Hirsinger and Steventon, 2017); 2) adjusting microscopy setups to produce high resolution images and low photo-toxicity (Garcia et al., 2011; McConnell et al., 2016); 3) customizing imaging software and hardware (Mc Dole et al., 2018); 4) post-processing of the acquired data, e.g. deconvolution (Sage, et al., 2017; Preibisch et al., 2014) and registration (Preibisch et al., 2010); 5) assembling efficient processing and analysis computing hardware (Roger et al., 2016); 6) segmenting and/or tracking cells in 3D over time (Schmidt, et al. 2018; Weigert et al., 2020; Tinevez et al., 2017); and finally, writing bespoke code to analyze the dynamics of the tracked cells (de Medeiros et al., 2021; Zhisong et al., 2020). Thus, without standardized pipelines in place, analysis of spatiotemporal cell dynamics can be a daunting task. Keeping increased accessibility as a central goal, here we provide a user-friendly cell-tracking pipeline accompanied by guidance, open-source code and novel analysis software.

We first give an overview of our pipeline, explaining the goals and concepts of each module, as well as discussing application and limitations. In Materials, we provide parameters and settings optimized to study the segmentation clock in zebrafish (Rohde et al., 2021). Step-by-step details for each module are given in the Procedures, which also includes guidance on modifying parameters for other samples.

### Pipeline Overview, Application and Limitations

The pipeline has 4 main modules (Figure 1): 1) a time-lapse of a live sample is acquired; 2) the time-lapse is processed to facilitate data handling and further analysis; 3) cells are detected as spots and tracked within the time-lapse and 4) spatiotemporal features are extracted from the cell tracks and analyzed.

**Figure 1.**
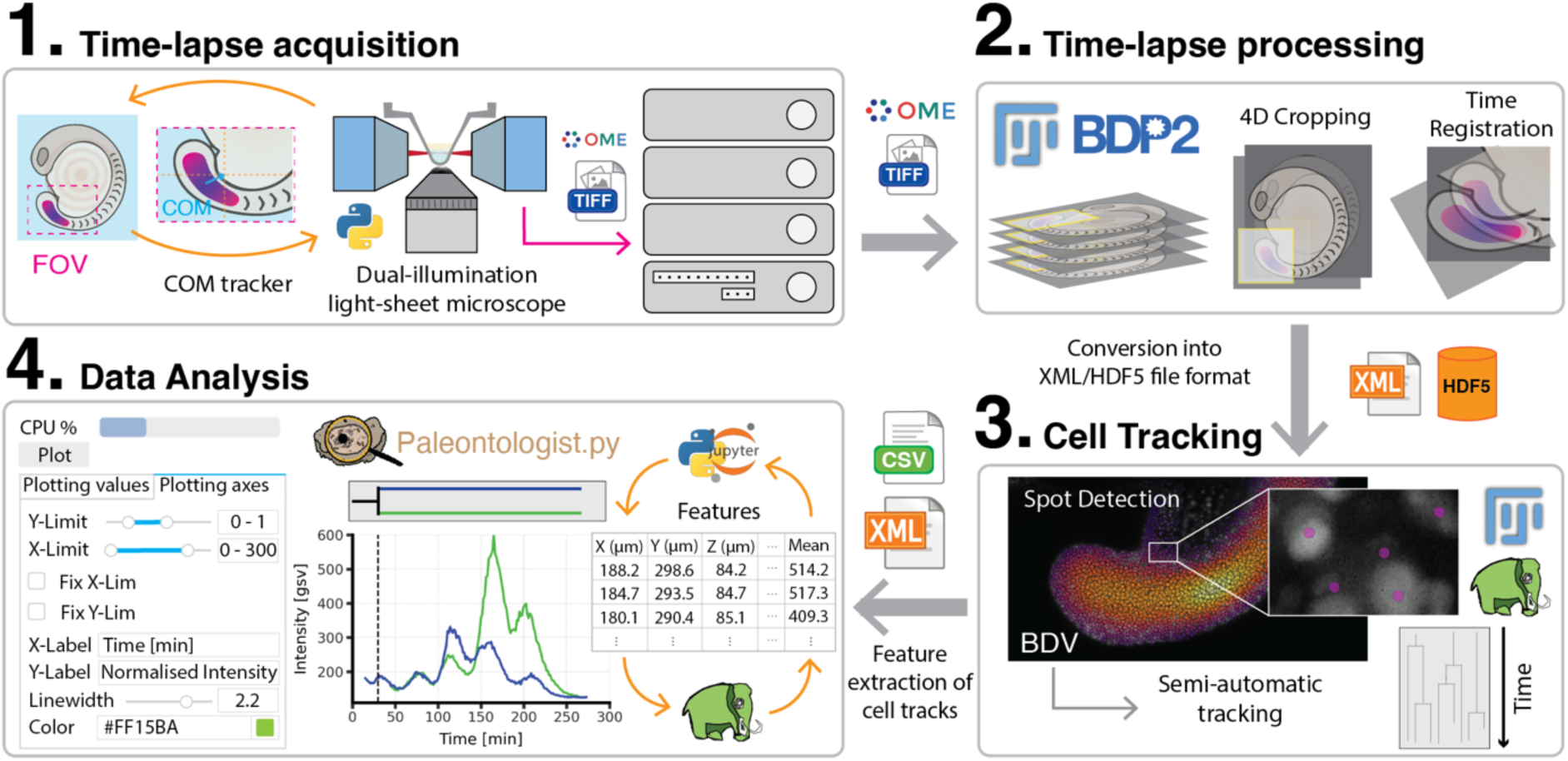
Cell tracking pipeline overview. **1)** Cellular resolution time-lapse acquisition using, for example, light-sheet microscopy. To keep the region of interest inside the field of view (FOV), Python-based Centre Of Mass (COM) tracking is implemented. Resulting OME-TIFF data is saved into a centralized workstation. **2)** The 5D time-lapse data is processed by cropping, then drift correction and time-registration to facilitate later cell tracking is applied. Processed time-lapses are saved in XML-HDF5 file format to allow interaction between the visualization and analysis tools. **3)** Tracking is done using Mastodon, a Fiji plugin. Mastodon outputs all the features from the cell tracks (XYZ cell coordinates, intensities, velocities, etc.) as an XML or CSV file. **4)** Data analysis of the features is made easy in Paleontologist, a modular python package that was built in our lab to interactively analyze tracked cells and output publication quality figures. Features from Mastodon can be iteratively plotted and edited in Paleontologist, then re-checked in Mastodon for cell visualization in the context of the embryo.

Here we demonstrate the step-by-step application of our pipeline using the example of our work following individual cells throughout segmentation of the developing zebrafish embryo (Rohde et al., 2021). The segmentation clock is a multi-cellular patterning system that translates the rhythm of cellular genetic oscillations into the successive and periodic formation of blocks of tissue in the trunk and tail called somites. Clock activity produces tissue-level waves of gene expression in presomitic mesoderm (PSM) that travel anteriorly until arrest at the position of the newly forming somite (Aulehla et al., 2008; Delaune et al., 2012; Masamizu et al., 2006; Palmeirim et al., 1997; Soroldoni et al., 2014). Historically, rapid cellular-level clock oscillations and ongoing tissue morphogenesis have made it difficult to describe the full picture of cellular dynamics underlying the clock pattern in zebrafish and other model systems (Delaune et al., 2012; Morelli et al., 2009; Shih et al., 2015; Yoshioka-Kobayashi et al., 2020). Consequently, the segmentation clock is a challenging system for the optimization of live imaging, visualization and analysis. In creating the cell tracking pipeline our motivation was thus two-fold, first to directly answer questions about cellular clock dynamics, and second, to standardize a pipeline that makes this level of analysis accessible to a broader range of researchers and model systems.

#### 1. Time-Lapse Acquisition

Cell-tracking in our system relied on a fluorescent nuclear marker, however the pipeline could also be adapted to track intra- or inter-cellular spot-like structures, for example tracking centrioles (Erpf et al., 2020). Feasibility was in large part determined by the quality of the acquired data; thus, our first step was always to consider sample-dependent limitations including constraint-free mounting of the live sample, photo-bleaching, photo-toxicity, and spatiotemporal resolution relative to the dynamics of interest. Each experimental system presents unique requirements, thus troubleshooting guidance is offered in the Procedures.

Depending on the sample size and microscopy hardware, the field of view (FOV) required to image at cellular resolution could fail to cover the entire region of interest (ROI). Particularly challenging is that the ROI itself could simply move out of the FOV due to growth and morphogenesis, a limitation we faced while imaging the extending tail of the zebrafish. Fortunately, we resolved FOV problems using short scripts of code to communicate with the microscope software controlling the camera and stage movement. For example, to enable long-term imaging of the segmentation clock, we designed a center of mass tracker that kept the fluorescent clock signal inside the FOV as the embryo extended its tail. Our tracking script, included here (see step by step in Fig S1), can easily be translated into other microscope systems that allow custom scripts. The automation enabled by such scripts will further advance adaptive imaging (Roger et al., 2016). For example, the custom software TipTracker is used to automatically track diverse moving objects on various microscope setups (Von Wangenheim et al., 2017).

#### 2. Processing

Following the acquisition of a TB-sized time-lapse movie, even the initial visualization could be problematic without the correct software and hardware tools due to the limitations of a standard computer’s RAM. To guide users over this hurdle, we established pre-processing steps to convert the time-lapse into a manageable, ready-to-be tracked format. These steps, which we detail in the Procedures section, included cropping to reduce size in all dimensions XYZT, conversion to HDF5 files, and time registration to reduce sample movement and drift.

Our pipeline thus allows broader users to consolidate and smooth the workflow through pre-processing steps that have previously been published as stand-alone operations (cropping, registration, chromatic aberration corrections), either custom built for a specific project (e.g., tracking of all the cells in a developing mouse embryo, McDole et al., 2018) or available as a commercial product (Imaris (Bitplane), Arivis). The open-source and commercial tools we recommend here for pre-processing have already been tested in multiple systems, and include a user-friendly interface (Fiji as open source; Imaris (Bitplane), Arivis as commercial). To facilitate the transfer and storage of the TB-sized datasets, we recommend data compression systems (Lempel-Ziv-Welch (LZW) and Deflate compression as open source; Jetraw (Dotphoton) as commercial).

#### 3. Cell Tracking

Pre-processing produced files that were easily opened and viewed in the cell-tracking tool Mastodon – a large-scale tracking and track-editing framework for large, multi-view images (https://github.com/mastodon-sc/mastodon). Although automation of cell tracking within Mastodon was limited by time-lapse quality, we cover in Procedures (step 14) how tracking parameters can be tuned for particular spot size, signal intensity, cell density, movement, etc.. Manual tracking and editing were also user-friendly. Features including mean intensity, XYZ coordinates, number of links, and velocities were extracted from the tracks and saved in CSV (comma separated values) files for later analysis.

#### 4. Data Analysis

To explore features extracted from the cell tracks we created Paleontologist (https://github.com/bercowskya/paleontologist), a novel open-source analytical tool that requires no coding experience, but allows custom scripting. Paleontologist was designed to interactively aid in quantitative and qualitative analysis of spatiotemporal features for single or multiple cell tracks of interest. Importantly, Paleontologist facilitated moving back and forth between Paleontologist and Mastodon to investigate, correct and refine cells or groups of cells of interest. After using the data exploration interface, we used Paleontologist to output publication-quality figures.

We were motivated to develop Paleontologist due to the limited tools available to analyze spatiotemporal dynamics, especially those from individual cell tracks. Louveaux and Rochette developed an R package mamut2r (https://marionlouveaux.github.io/mamut2r/), which imports and visualizes xml files from MaMuT, a Fiji Plugin precursor to Mastodon (Carsten et al., 2018). However, besides custom-built scripts for specific purposes, no open-source tool existed to perform similar tasks for Mastodon. Mastodon outputs large CSV files that include the necessary information to reconstruct cell tracks, however the reconstruction process was often challenging in the presence of cell division. Paleontologist solved these issues, returning arrays of reconstructed tracks that include an ID for cell division to keep track of daughter cells. Moreover, spatiotemporal analysis was complicated due to the need to consider data pre-processing. For example, if registration was performed, then coordinates provided by Mastodon would also need to be registered. Paleontologist solved this by allowing the registration to be undone if needed.

## Materials

### 1. Zebrafish

Transgenic (Tg) fish were maintained according to standard procedures in École Polytechnique Fédérale de Lausanne (EPFL, Lausanne, CH). Using natural pairwise spawning, we obtained double Tg embryos heterozygous for a real-time segmentation clock reporter *Tg(her1:her1-yfp)* (Soroldoni et al., 2014) and the nuclear marker *Tg(Xla.Eef1a1:H2BmCherry)* (Recher et al., 2013). Embryos were incubated in facility water at 28.5°C until shield stage, then 19.5°C until the 8 to 10 somite stage, after which they were returned to 28.5°C until imaging at the 15-somite stage. Embryos were dechorionated manually prior to imaging, then immersed in facility water with 0.02% Tricaine (Sigma) during imaging to avoid muscle twitching.

### 2. Microscope

We used a LightSheet Microscope LS1 (Viventis Microscopy Sárl, Switzerland) with the following configuration: Andor Zyla 4.2 sCOMS camera; 515 nm laser to image YFP; 561 nm laser to image mCherry; CFI75 Apochromat 25X, NA 1.1 detection objective (Nikon); scanned gaussian beam light sheet with thickness (FWHM) of 2.2 µm.

### 3. Imaging

#### 3.1 Mounting

Whole embryos were mounted in an imaging chamber that reliably holds them in a lateral orientation, ideal for illuminating the segmenting trunk and tail. To make our molds, we first glued a thin membrane to the bottom of a 3D-printed chamber to make a trough (Viventis Microscopy Sárl, Switzerland). 2% LMP Agarose (Sigma, in E3 medium) was then added gradually to the trough along with a 3D-printed counter mold of 5 to 10 small protruding semi-circles (750 µm in diameter) such that depressions were created to hold the embryo’s yolk while allowing unhindered extension of the tail and body (Herrgen L., Schröter C., Bajard L., Oates A.C., 2009) (Figure 2B). Embryos were added to the chamber after removing the mold and filling the trough with facility water plus 0.02% tricaine (Sigma). Our region of interest, the trunk and tail, lay flat in a lateral view along the thin agarose surface (Figure 2B). Temperature was kept at 28.5°C using a recirculating air heating system (Cube 2, Life Imaging Services, Switzerland).

**Figure 2.**
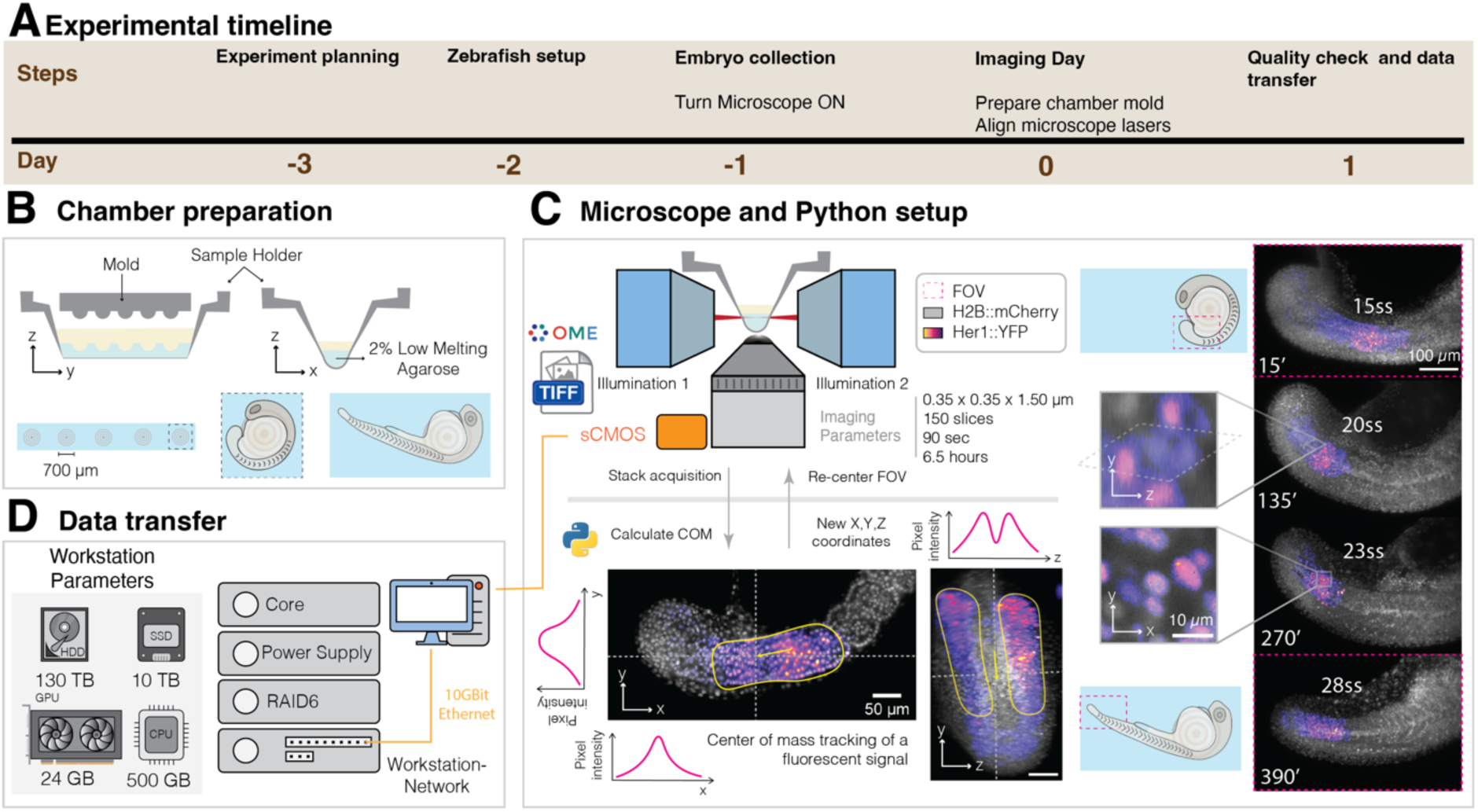
Time-lapse acquisition of zebrafish embryonic segmentation. **A)** Timeline to acquire a time-lapse of a zebrafish embryo. **B)** Preparation of the imaging chamber and mounting the embryo. A 3D-printed sample holder was glued to a transparent filament sheet, creating a trough. Low-melting point agarose was added to the trough, then a 3D printed mold was used to create depressions in which the yolk of the embryo sits while the tail freely extends. **C)** We used a dual-illumination light-sheet microscope to acquire 5D time-lapse movies. Data was saved as an OME-TIFF file. During acquisition, the center of mass (COM) of the signal of interest (Magenta), was tracked to instruct the microscope to re-center the field of view (FOV) in XYZ. **D)** Acquired data was transferred from the imaging computer to an image processing station.

#### 3.2 Imaging parameters

To track cells, we relied on a non-oscillating nuclear marker *Tg(Xla.Eef1a1:H2BmCherry)* (Recher et al., 2013). Cells in the segmenting region of the zebrafish embryo have a nucleus of 7-10 µm, requiring z-planes every 1.5 µm to produce spatial resolution suitable for tracking. We took stacks of 150 z-planes to span the depth of one entire side (right or left) of the bilaterally segmenting tissue at 15 somite stage and older. Younger embryos required more z-planes to compensate for greater depth of the segmenting tissue in the lateral orientation. To reliably follow individual cells semi-automatically, we needed to acquire images every 90-120 seconds due to cell movement and mixing in the segmenting tissue.

Our time-lapse movies ran for at least 6 hours (240 time points), with 90 sec intervals, a bit depth of 16, X-Y dimensions of 2048×2048 pixels, two channels, and 150 planes. A movie with these parameters has the following raw size:

Time Lapse movie size = [240 × 16 × 2048 × 2048 × 2 × 150] bits × 1 byte/8 bites = 6 × 10^11 bytes = 600 GB

#### 3.3 Center of mass tracking

To keep the segmenting tissue in the FOV, we automatically tracked the center of mass (COM) of the Her1-YFP signal while acquiring the time-lapse (Figure 2C). COM detection was performed using a python environment that directly communicates updated coordinates to the microscope control system. To find the COM in our channel of interest, YFP, we use a cropped region (Figure S1A) of a single timepoint that had been XYZ max-projected in the YFP channel (Figure S1B). These projections were processed using a median filter and a gaussian blur to smooth the signal, resulting in a filtered max projection that we binarized using an Otsu thresholding method (Figure S1C). COM was then calculated using this binary mask, and an offset value was produced corresponding to the XYZ distance that the FOV needs to shift when re-centering (Figure S1D). To prevent abrupt shifts, we set the movement to be maximum of 5 µm per interval (Figure S1E). The following additional parameters could be adjusted: filter size applied (Gaussian and Median); use of the entire field of view or a cropped section (in XYZ); start of tracking while imaging; and COM as a binary mask, in which the center of pixels was used, or an intensity mask, in which the brightest area acted as the COM.

#### 3.4 Data compression

The resulting four-dimensional (XYZT) for each channel was saved as an OME-TIFF (Leigh et al., 2017; Besson et al., 2019) (Figure 2C), a standardized format that is read by most open-source and commercial software. To make data handling easier during moving, processing and storage, we recommend compressing the data during acquisition so that the output is a compressed OME-TIFF file, which is easily read in Fiji. LZW (Lempel–Ziv–Welch) is an open-source universal lossless data compression algorithm that is easy to implement. In the case of our time-lapse movies, we obtained a reduction factor of 1:2, and could compress the data during acquisition if the imaging parameters permitted this in terms of speed. If the compression speed was too low to run during acquisition, or a better reduction factor was desired, a commercial solution, Jetraw (Dotphoton SA) allowed for a 1:8 compression ratio at acquisition speed.

#### 3.5 Quality Check

To avoid saving poor quality time-lapse data, e.g., a movie in which the sample degrades, we performed a quality check in parallel to acquisition by saving a maximum-intensity projection for each timepoint as a tiff file (Figure S2) (see Table 1). The max projections were viewed during acquisition (using Fiji) without the memory and speed problems that would otherwise be caused if trying to view the whole stack. Quality was checked by assessing tissue structure, signal to noise ratio and photo-bleaching over time.

**Table 1.**
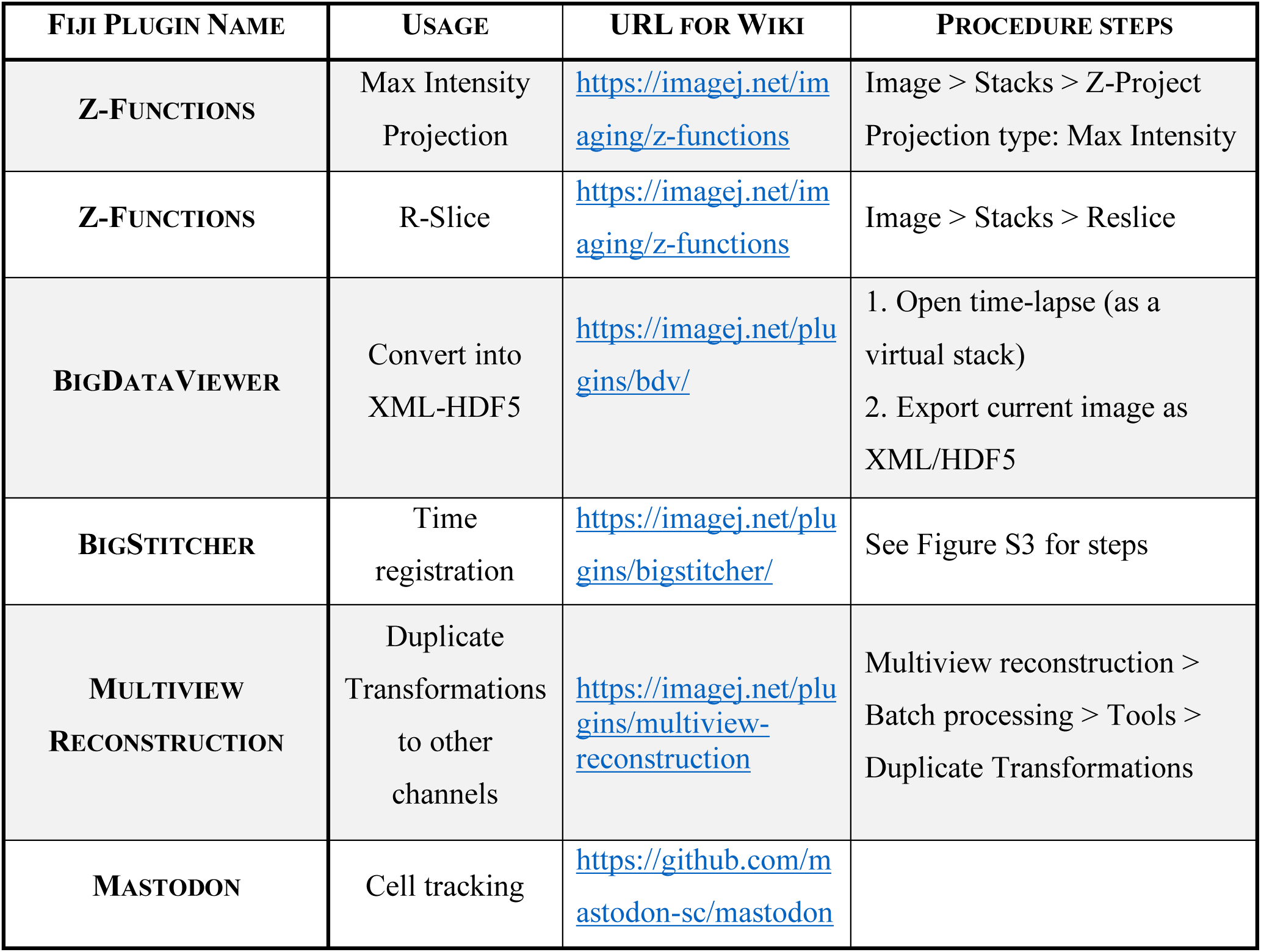
Fiji Plugins name, usage, link to wiki and procedure steps.

#### 3.6 Time registration

To stabilize the time-lapse images in the time dimension to improve the cell tracking, we registered the 3D volumes from all timepoints using the first time-point as the frame of reference. Registration parameters were tuned according to the sample and imaging parameters. We show the steps and the parameters used in Figure S3, which can serve as a starting point.

### 4. Computing Hardware

Image processing and data handling was done using a HIVE (Acquifer Imaging), a powerful centralized workstation for big-data storage and high-performance computing. Our HIVE is equipped with a 24 GB Graphics Processing Unit (GPU), 500 GB of Random-access memory (RAM), 10 TB Solid-state drive (SSD) (RAID5) and 130 TB Hard Drive Disk (HDD) (RAID6). Data was stored in SSD while processing, then moved to the HDD for short-term storage. The microscope computer where the data was saved during acquisition was connected through a 10 Gbit cable to the HIVE to allow rapid transfer.

### 5. Software

Our main software platform was Fiji (Schindelin et al., 2012), which can be installed in Windows, Max OSX and Linux operating systems. The core functionality of Fiji was extended using plugins specified in our protocol. Fiji and its plugins can be found along with installation instructions at the imagej.net website.

Paleontologist runs on Python 3.6 or above and installation instructions are on the GitHub webpage (https://github.com/bercowskya/paleontologist). Install Anaconda Distribution (Anaconda Inc, 2020) to include interactivity and the user interface.

#### Procedure

##### Sample preparation and Image Acquisition

1. REQUIRED: Image the sample on a light-sheet or confocal microscope with resolution parameters that enable cell tracking (our parameters are shown in Figure 2C). Cell tracking works best in time-lapses acquired at high temporal and spatial resolution relative to cell movement/mixing. A rule of thumb to follow is that if the cell cannot be followed by eye, most likely the automatic tracking will not be successful. TROUBLESHOOTING: Explore the range of spatiotemporal resolution in which tracking is feasible by acquiring short time-lapses using various parameters, then view as described in step 2 to check if individual nuclei can be followed by eye. Balance this acquisition rate against potential photo-toxicity and the projected file size over the time interval of interest.
2. OPTIONAL: Use a COM tracker (or other method that enables automated detection of the ROI) to keep the ROI in the FOV (Figure 2C, S1).
3. OPTIONAL: Data compression during or after acquisition is highly recommended (LZW, Jetraw).

##### Quality check of time-lapses and data transfer

4. REQUIRED: Download the free, open-source image processing software Fiji (https://imagej.net/software/fiji/#downloads)(Schindelin 2012). Note that there are commercial and other open-source solutions for big image data inspection including Imaris (Oxford Instruments), Arivis Vision 4D (Arivis AG), Vaa3D (Bria, et al., 2015) and TDat (Li, et al., 2017). Here, the focus is on processing and tracking plugins using Fiji.
5. REQUIRED: Generate a maximum-intensity projection of the time-lapse (Table 1) using image processing software of choice, then check the following to assess time-lapse quality:

a. Visually check for photo-bleaching of the signal over time. A severe intensity decay could obscure the dynamics of interest, as well as interrupt cell tracking (Figure S2A). TROUBLESHOOTING: Either alter the imaging parameters when possible (reduce exposure time, laser intensity, frame rate) or correct using a tool such as correction for photo-bleaching from Miura, 2021.
b. Using the re-slice tool from Fiji (Table 1), check whether the z-resolution is high enough so that the cells look like spot-like structures (Figure S2B). If the resolution is too low, the cells will appear almost like lines and the spot detection during cell tracking will not work. TROUBLESHOOTING: Alter imaging parameters by reducing the pixel size in the z-axis.
6. REQUIRED: Transfer the checked time-lapse to an image processing station. The specifications vary according to the size of the data and the desired waiting time between processes. Because all of the tools we propose here can handle big data, increasing RAM or GPU availability will only improve the speed of the processing and the interactivity during the cell tracking. BigDataViewer (BDV) adapts the size of the cache to the available memory (Pietzsch et al., 2015) and BigDataProcessor2 uses lazy loading and processing (Tischer, et al., 2021).

##### Data conversion

7. OPTIONAL: Crop the time-lapse in XYZT to reduce file size (Figure 3B) using BigDataProcessor2 (BDP2, Tischer, et al., 2021), a Fiji plugin for processing n-dimensional big data images. Install by activating the BigDataProcessor Fiji update, then access using the graphical user interface or a Fiji macro. The time-lapse OME-TIFF is loaded and displayed using BigDataViewer (BDV, Pietzsch et al., 2015), which allows efficient lazy loading of raw data such that all processing steps are applied and then re-saved only once. When cropping, confirm that the sample remains inside the bounding cropping box by checking the initial, intermediate and last timepoints.
8. OPTIONAL: Detect chromatic shifts by looking for small XYZ shifts in normally overlapping signals (e.g., a nuclear marker and nuclear localized signal). Correction for shifts can be applied uniformly throughout the time-lapse in BDP2.
9. REQUIRED: Convert the time-lapse to HDF5 format (The HDF Group, 1997-2019) using BigDataViewer (Pietzsch, Tobias, et al., 2015). The HDF5 is associated with an XML file containing the metadata and all future registrations, etc. applied to the data. The XML-HDF5 should be stored in an SSD for fast reading and writing operations. When the conversion starts, the HDF5 and the XML are automatically created at the same time. All subsequent steps use HDF5/XML files. TROUBLESHOOTING: The XML file and the HDF5 need to be in the same folder because the XML file has the path of the HDF5 used when it was created. Therefore, if the XML and HDF5 are separated, a reading error will appear when trying to open the data.

**Figure 3.**
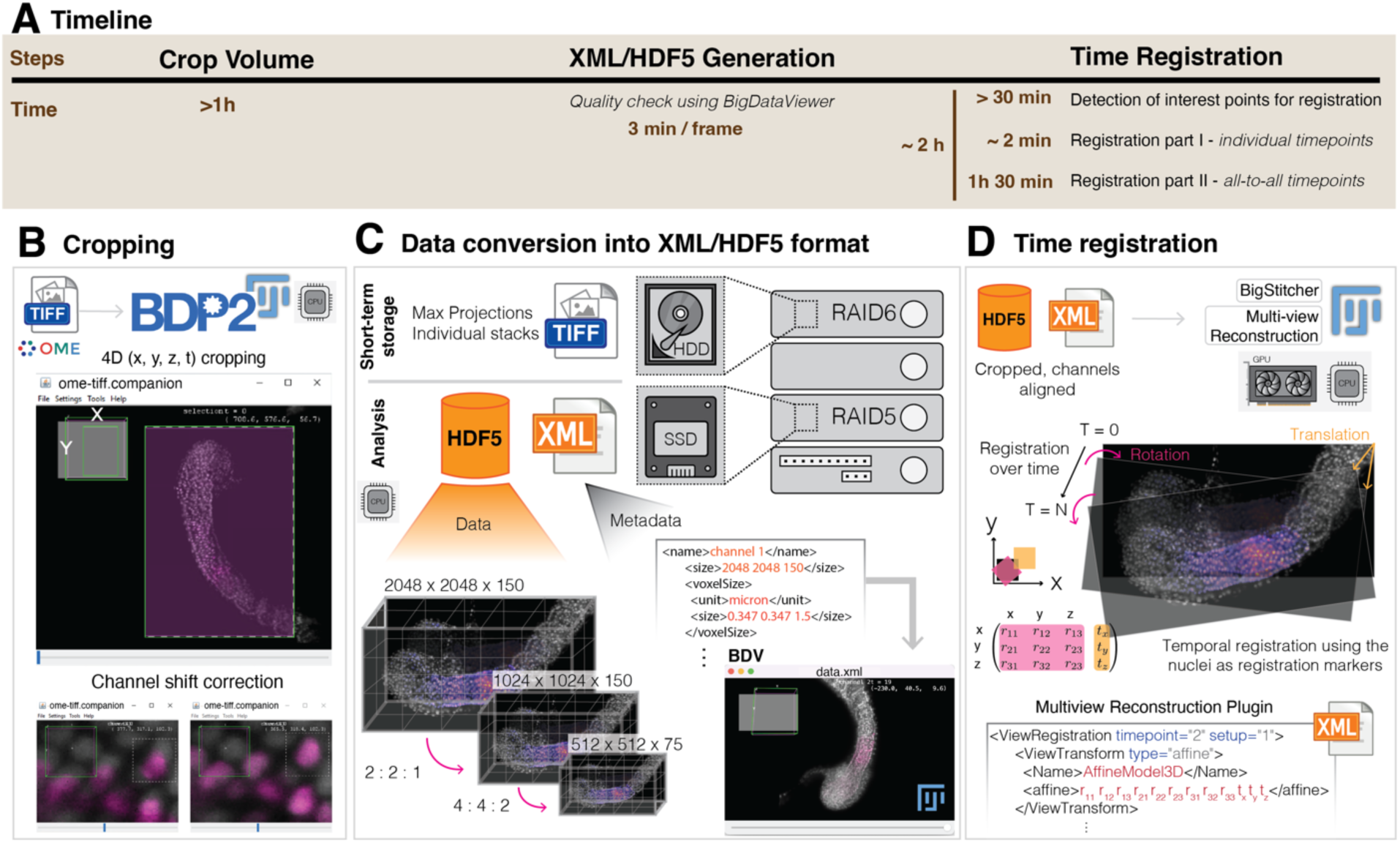
Pre-processing the Time-lapse. **A)** Timeline of pre-processing using the specified memory parameters from Figure 2B. **B)** Cropping and channel shift correction was performed using BigDataProcessor2. After loading the OME-TIFF files, the user interface offers a choice of transformations to apply (affine transformed viewing, cropping, binning, bit-depth conversion, drift correction and channel alignment). **C)** The transformed and cropped data was saved into XML-HDF5 file format in the SSD to speed future read-write processes, while the OME-TIFF raw data was stored in HDD as backup. HDF5 data is organized in a pyramidal structure that enables interactivity when opened in BigDataViewer. **D)** To correct for embryo movement or drift, time registration was applied using the nuclei as registration markers. This step could be performed in CPU or GPU, with the resulting registration matrices for each timepoint stored in the companion XML file.

##### Time registration

10. REQUIRED: During extended time-lapse imaging, the sample might drift due to growth or technical issues. This can make the cell tracking harder or even impossible. To compensate for this drift, the nuclei are used as markers to register individual time-points to each other. Select a timepoint, usually the first, then use it as a reference. To perform time registration, use Fiji Plugins Multiview-Reconstruction (Preibisch et al., 2010) or BigStitcher (Hörl et al., 2019) as both are compatible with the XML-HDF5 file format. An alternative to Fiji would be Elastix (Klein et al., 2009; Shamonin et al., 2014), a toolbox for intensity-based medical image registration. If the imaging setup allows it, for instance when performing multi-view imaging, beads can be added and then used as registration markers (Preibisch et al., 2010).
TROUBLESHOOTING: The registrations are not actually applied to the data, but rather the matrices applied are saved for each timepoint in the XML (Figure 3D). This is useful because backup XML files (saved as ∼.xml) are created in the process so that in case the registration fails, you can go back to the unregistered XML file and try various parameters without having to re-convert the data into XML-HDF5.

a. Detect nuclei using the feature “Detect interest points”. Use the detection method “Difference of Gaussian (DoG)”. Two parameters need to be defined for detection of interest points, 1) an intensity threshold, and 2) a radius. These parameters can be tested on a single time-point before running the detection for the whole time-lapse. If the nuclear signal varies over time (for example in the case of photobleaching), tune the detection parameters using a time-point where the signal is weak.
b. Perform a first round of registration using the method “Fast description based” (rotation invariant) registration in which timepoints are registered individually. Moreover, all the views are compared to each other and the first time-point is fixed so that the rest of the time-points can map back to it using the translational invariant model. Use the transformation model rigid affine. Figure S3A shows the parameters applied to segmentation clock time-lapses, which can be used as a starting point to tune parameters for other systems.
c. Perform a second round of registration using the method “Fast description based” translational invariant. In this case, because all timepoints are registered, group-wise optimization by reasonable global optimization to “all-to-all” time-points with range needs to be performed. As before, the first view is fixed and the rest are mapped back using a translational model. Parameters used for the segmentation clock time-lapses (Figure S3B) will require fine tuning for each independent dataset.
d. Using the Fiji plugin “Multiview reconstruction”, duplicate the transformation obtained using BigStitcher for the “nuclear marker” channel to the other channels (Multiview Reconstruction > Batch processing > Tools > Duplicate transformations).
e. Apply transformation of “One channel to other channels”. TROUBLESHOOTING: Registration usually fails due to the poor alignment of the registration markers (nuclei) over time. This can be corrected by improving the temporal resolution. Bright objects in the FOV that are not within the sample, for example lint or debris, can also disrupt registration. TROUBLESHOOTING: Automatic registration can sometimes fail because of the sample ‘jumping’ or rotating significantly in a few timepoints during the acquisition (the embryo might fall on the side and adopt a new equilibrium position). The resulting discontinuities may prevent automatic tracking. It is possible to correct big discontinuities manually with a set of tools from the BigDataViewer-Playground library (https://imagej.net/plugins/bdv/playground/bdv-playground-manual-registration).

##### Spot detection and cell tracking

11. REQUIRED: The dataset is now ready for cell tracking using the Fiji plugin “Mastodon”. Set up three BDV windows with orthogonal views. Synchronize the windows by locking each of them on view 1 by clicking on the first lock (Figure 4A). Select the channel corresponding to the nuclear marker, which will be used for tracking.
12. Set up the TrackScheme window and synchronize it with the BDV windows by locking each of them on view 1 by clicking on the first lock.
13. Familiarize yourself with the actions and their corresponding keyboard shortcuts. They can be found, and modified, in Mastodon > File > Preferences > Keymap.
14. Set up semi-automatic tracking (Mastodon > Plugins > Tracking > Configure semi-automatic tracker…) with the following parameters (information about the parameters is displayed by placing the cursor over a given parameter):

a. Setup ID: Select the channel corresponding to the nuclear marker. This is the channel that will be used for tracking.
b. Quality factor: This parameter depends on the dataset and needs to be empirically determined. A value of 0.5 is a reasonable starting point.
c. Distance factor: This parameter depends on the dataset, especially on how much cells move between time frames, and needs to be empirically determined. A value of 1.5 is a reasonable starting point.
d. N time-points: This parameter specifies how many time-points can be processed at most. It does not affect the quality of tracking itself and can therefore be set at a large number (e.g., 40).
e. Tracking direction selection: Forward- and back-tracking in time can be performed.
f. Untick “Allow linking to an existing spot” if performing semi-automatic tracking to prevent the new track starting from a track that was previously curated.
g. Tick “Run detection if existing spot cannot be found”
15. Place the cursor on one cell of interest and hit “A” to add a new Spot.
16. Adjust the size of the Spot to make it fit the nucleus by making the Spot smaller (“Q”) or bigger (“E”).
17. Click on the cell of interest, place the cursor inside the Spot added in step 14 and start semi-automatic tracking (Mastodon > Plugins > Tracking > Semi-automatic tracking or Ctrl + T). TROUBLESHOOTING: If semi-automatic tracking fails, the first step is to adjust the tracking parameters in Mastodon. Second, to optimize trackability, reiterate tracking attempts in timelapses with improved registration and different imaging settings.
18. Curate the track by visual inspection. Check that the cell of interest is followed through the entire track. Portions of a track can be deleted onwards from the timeframe where an error is made. OPTIONAL: tag the track with a label, for example “checked”.
19. Save the Mastodon project regularly to avoid losing tracking data in case of software crash (“Save” button in the Mastodon window or Mastodon > File > Save project)
20. Repeat steps 17 to 19 until the cell of interest has been tracked for the desired duration.
21. Repeat steps 15 to 20 for each cell that needs to be tracked.
22. Compute the features of interest (“compute features” button in the Mastodon window).
23. Generate a results table (“table” button in the Mastodon window).
24. Export the table as a CSV file (Table window > File > Export to CSV).
25. To use Paleontologist, load the CSV and the XML files. To see the details, visit https://github.com/bercowskya/paleontologist.

**Figure 4.**
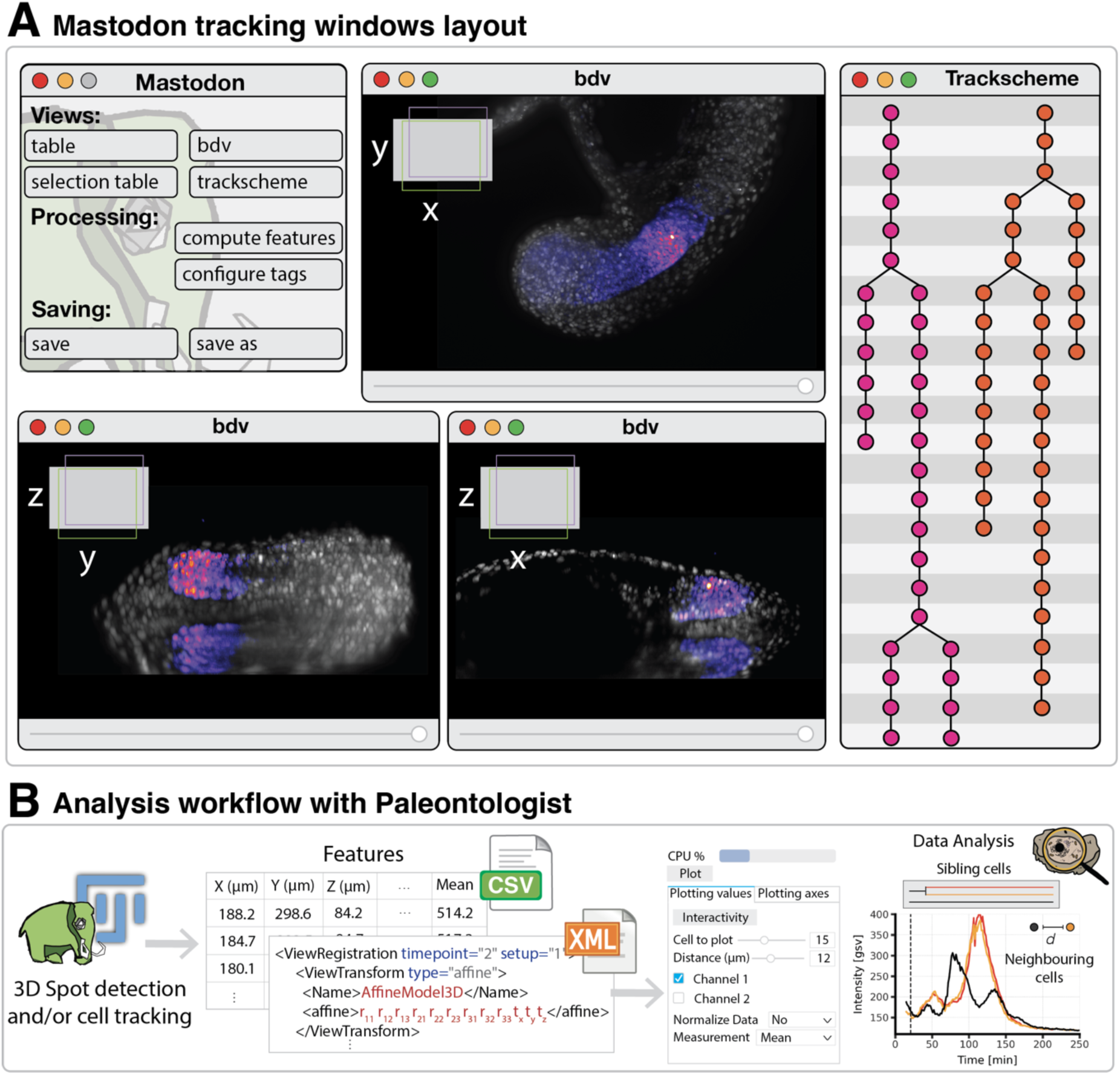
Mastodon and Paleontologist workflow: **A)** Mastodon allows interactive tracking in 3D. Cells can be selected and tracked, manually or semi-automatically, within multiple BigDataViewer (bdv) windows. An accompanying trackscheme facilitates manual editing and tagging of individual cell tracks. **B)** Tracking data is exported from Mastodon as CSV and XML files containing features including cell coordinates, cell intensity, number of links per spot (to later reconstruct divisions) and frames. Paleontologist reads these files, then allows both interactive analysis and editing for re-inspection of specific cell tracks in Mastodon. Here we show an example of the user interface to explore cell division and neighboring cells. Values to plot can be selected, including channel, individual cell(s), distance to neighboring cells, and measurement used to calculate the intensity (mean, median, etc.). Axes, plotted lines and labels are all tunable for optimum presentation of the data.

## Acknowledgements

We thank JiSoo Park, Pierre Osteil, Alexandre Mayran for testing the pipeline and generating feedback, J-Y Tinevez and T. Pietzsch for Mastodon assistance, EPFL’s fish facility and Bioimaging and Optics Platform, P. Strnad and A. Boni for imaging help.

## Supplementary Materials

Figures S1 to S3

