## Supplementary material for "But, what are the cells doing? Image Analysis pipeline to follow single cells in the zebrafish embryo" (from another version of this preprint: version 1, 2023-06-04)

**Affiliations:**


**This PDF file includes:**

Page 2-3: Figures S1 – S3

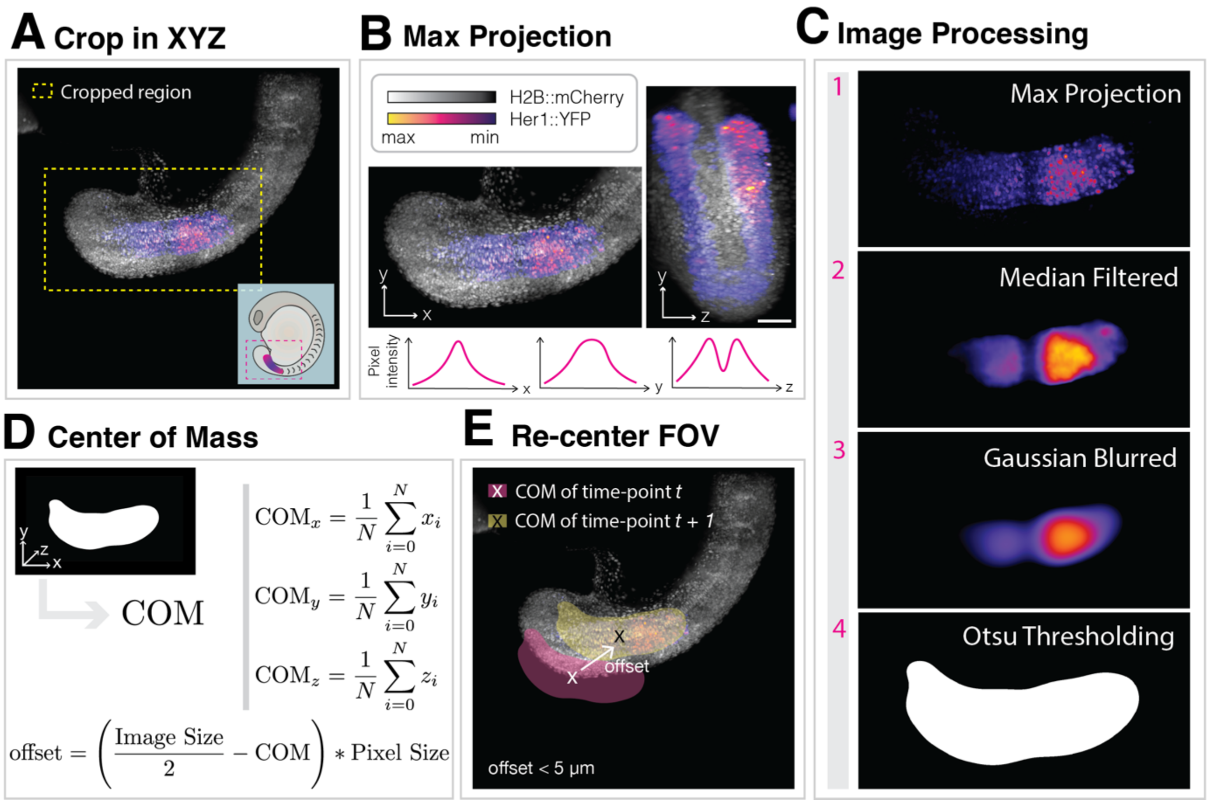


**Figure S1. Centre of Mass calculation to re-centre the field of view. A)** Crop the data in XYZ in order to restrict processing to the region of interest. **B)** Max project the cropped volume in the channel desired for COM tracking. **C)** Perform image processing steps to make it easier to obtain the COM. First, apply a median filter (filter size = 40 in this case), and a gaussian blur (σ = 40). Then apply an Otsu threshold to binarize the images. **D)** Calculate the centre of mass (COM) from the binary image. This is done by performing the sum of all the coordinates (xyz) where the binary mask is equal to 1 and dividing by the number of pixels where the mask is equal to 1. The offset is then calculated using the image size and pixel size. **E)** The offset, as long as it is not greater than 5 µm, is applied to the next time-point in order to re-centre the field of view.

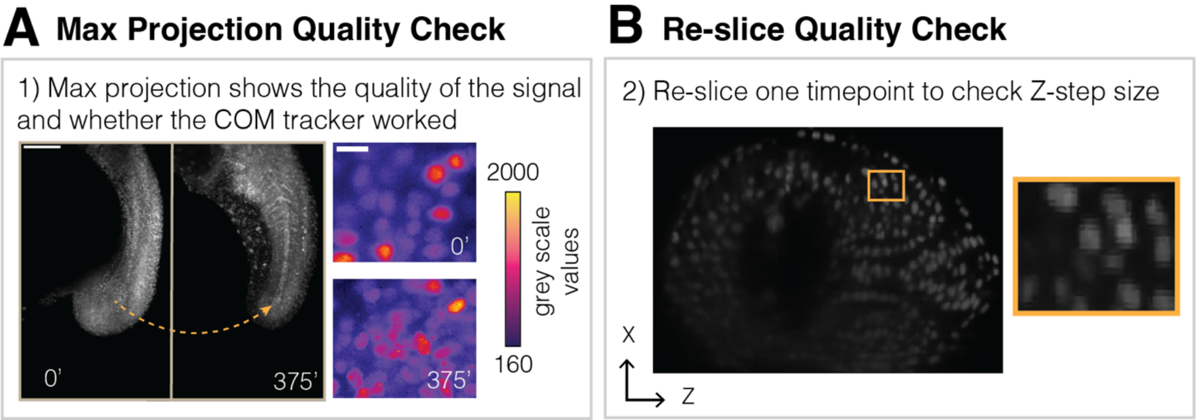


**Figure S2. Quality checks. A)** Once the acquisition is over, the first thing to check is whether the COM tracker worked by max projecting to see if the ROI remains in the FOV. Photo-bleaching and bleed-through can be detected over time by displaying a histogram in Fiji by pressing the letter "h" (https://imagej.nih.gov/ij/docs/menus/analyze.html#hist). Error bars displayed (left to right) are 100 µm and 10 µm. **B)** To observe whether the z-step size is sufficient to observe trackable blob-like structures, the re-slice option in Fiji can be used to change the view at a given timepoint. For example, these nuclei have a diameter of 10 µm and the z-resolution is 1.5 µm.

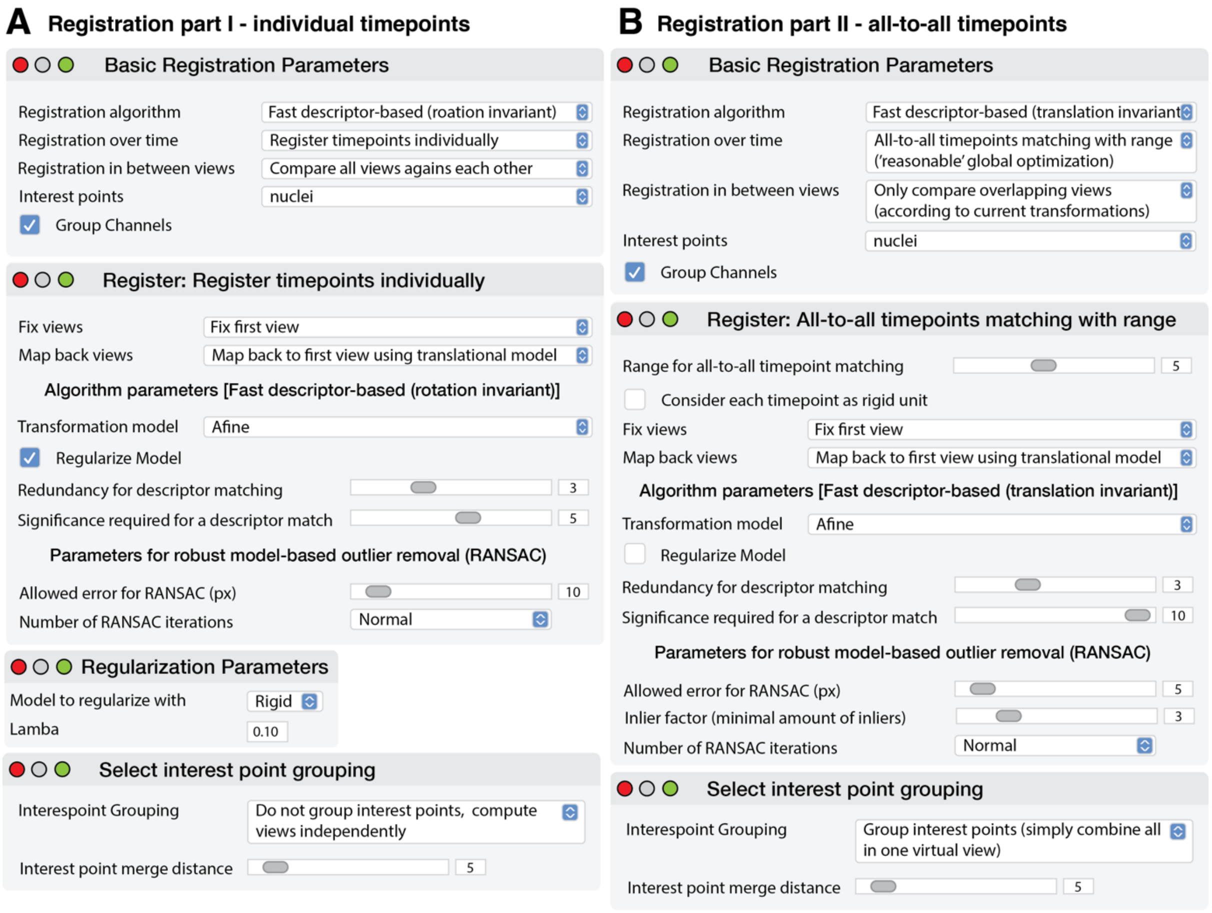


**Figure S3. Time Registration step-by-step. A)** Load the XML file and the user interface will show. First, register the timepoints individually using rotation invariant registration. **B)** As a second round of registration, reload the XML file and the user interface will reappear. This time, registration will be translation invariant and applied to all timepoints matching with range.
